# Do priority effects outweigh environmental filtering in a guild of dominant freshwater macroinvertebrates?

**DOI:** 10.1101/216267

**Authors:** Chelsea J. Little, Florian Altermatt

**Affiliations:** Department of Aquatic Ecology, Eawag: Swiss Federal Institute of Aquatic Science and Technology, Dübendorf, Switzerland; Department of Evolutionary Biology and Environmental Studies, University of Zürich, Zürich, Switzerland

**Keywords:** Amphipod, aquatic ecology, community assembly, competition, intermittent streams, land use, metacommunity, shredder, species distributions

## Abstract

Abiotic conditions have long been considered essential in structuring freshwater macroinvertebrate communities. Ecological drift, dispersal, and biotic interactions also structure communities, and although these mechanisms are more difficult to detect, they may be of equal importance in natural communities. Here, we conducted repeated surveys of locally-dominant amphipod species across ten naturally replicated stream catchments. We then used a hierarchical joint species distribution model to assess the influence of different drivers on species co-occurrences. The species had unique environmental requirements, but a distinct spatial structure in their distributions was unrelated to habitat. Species co-occurred much less frequently than predicted by their niches, which was surprising because laboratory and field evidence suggests they are capable of coexisting in equal densities. We suggest that niche preemption may limit their distribution and that a blocking effect determines which species colonizes and dominates a given stream catchment, thus resolving a long-standing conundrum in freshwater ecology.

---

> “It is reasonable to suppose that the fundamental niches of the two species overlap, but that within the overlap [the amphipod Gammarus] pulex is successful, while [the amphipod G.] deubeni with a greater tolerance of salinity has a refuge in brackish water…This case is as clear as one could want except that Hynes is unable to explain the absence of G. deubeni from various uninhabited favorable localities in the Isle of Man and elsewhere…These disconcerting empty spaces in the distribution of Gammarus may raise doubts as to the completeness of the picture.”
>
> — (Hutchinson 1957)

## Introduction

A central goal of ecology is to understand the factors determining the distribution of species, and the mechanisms of how these species are structured into communities. For instance, species distribution models based on environmental variables are commonly used to characterize species’ niches, and then to predict where they should be found. However, other processes are also important in determining species distributions, such as dispersal (Macarthur & Wilson 1967; Hubbell 2001; Leibold *et al.* 2004), interspecific interactions like competition (Wisz *et al.* 2013), and processes like order of arrival or “priority effects” (Drake 1991; Chase 2003; De Meester *et al.* 2016). Particularly at the local scale, these processes may in effect prevent the coexistence of species that are otherwise similarly suited to environmental conditions, and which do coexist at broader spatial scales (Thuiller *et al.* 2015). Compared to environmental variables, factors like order of arrival and dispersal limitation are not easy to detect or quantify in observational data, and so far have been largely neglected in species distribution models despite widespread recognition of their importance (Guisan & Thuiller 2005; D’Amen *et al.* 2017).

Amphipods are one of the lesser-known examples of organisms Hutchinson used in his seminal 1957 remarks positing the factors shaping species coexistence (see epigraph). While the examples on plankton or plant community coexistence are much more widely referred to, the relatively species-poor family of Gammaridae amphipods (Crustacea: Amphipoda) are an enigmatic case because the individual species are highly similar to one another ecologically, for instance using the same food resources, and are speculated to fill the same niches (MacNeil *et al.* 1997). Furthermore, in regions such as Europe and parts of Eurasia, they are the most dominant and important decomposers in freshwater ecosystems, thus playing a key role in ecosystems and food webs. In general, dominant species in a given community can structure communities and play an essential role in determining ecosystem function (Hillebrand *et al.* 2008). This is also true for amphipods, with greater dominance by the common central European species *Gammarus fossarum* associated with higher decomposition rates in streams (Dangles & Malmqvist 2004).

Because of such ecosystem-level effects, the distribution and potential coexistence of amphipod species are of particular interest. As noted by Hutchinson (1957) and others (for instance, Pinkster *et al.* 1970 called the amphipod species distributions a "problem"), mechanisms behind both these species’ commonly observed coexistence, but also the equally-common apparent exclusion of one by another, need clarification given that the species’ niches are assumed to be so similar.

In general, when a new species arrives from the regional species pool, there are three relevant outcomes in a community, assuming that the species’ abiotic requirements are met: (1) the new species cannot establish in the community; (2) it establishes and coexists with the other species; or (3) it establishes and replaces the previously-dominant species. The first case can occur when the species are functionally similar and niche space is not wide enough for both to coexist (niche preemption), or even when they are dissimilar but the previously-established species has modified and erased the niche of the new species (Fukami 2015). The second case can occur when species have different niches and is often considered desirable, as the presence of several functionally-different dominant species promotes stability in ecosystem functioning (Allan *et al.* 2011). The third case, meanwhile, is typical but not exclusive to invasive species. These three cases illustrate that even when there is the opportunity for multiple functionally-similar species to coexist, they not always do so. Furthermore, the “final” outcome of species interactions after a new species’ arrival depends on the temporal and spatial scale being considered. Coexistence in the short term may lead to species replacement (succession) over a longer time frame. Likewise, very small habitat patches may show a mosaic-like pattern of dominant species, while considering larger habitat patches may obscure this spatial segregation and indicate that species are coexisting.

To identify the mechanisms governing the distribution and coexistence of freshwater amphipods, we surveyed 121 stream reaches distributed in ten headwater stream catchments in Eastern Switzerland, sampling throughout the network topology of main and side stems, and capturing temporal dynamics by visiting each stream seasonally for one year. Previous work indicated that five amphipod species were present in the downstream lake, and three of these species consistently occupy the tributary catchments (Altermatt *et al.* 2016). These omnivorous species are comparable in size and functionally similar. They can move kilometers or tens of kilometers per year when expanding their ranges (Dick et al. 1994; Bollache et al. 2004; Piscart et al. 2008), and no genetic differentiation has been observed at the scale of the stream networks in this study, indicating unrestricted dispersal (Altermatt et al. 2016; note that there is genetic differentiation at larger scales, e.g. Westram et al. 2013). Thus, we assumed that the distribution of these different species across our study catchments should be driven only by niche differences with respect to abiotic conditions, biotic interactions, and/or stochastic processes, rather than dispersal limitation. We hypothesized that:

a. Species richness would be invariable throughout the sampling region, but with different species or combinations of species in different locations comprising this diversity; *or*,
b. Species richness would be higher at downstream points near the lake outlet. We would expect such a pattern both due to these points being in closest proximity to the regional species pool and due to typical diversity patterns found in river networks (Muneepeerakul *et al.* 2008); *and*,
c. Species would have individual niches and habitat preferences. This has been demonstrated for amphipod species in lakes (Hesselschwerdt *et al.* 2008) and larger rivers (Kley & Maier 2005), and we expected that this niche partitioning would explain why species were found in different locations. For example, such environmental requirements would lead to coexistence in complex habitats and/or to spatial segregation of species into non-overlapping patches within a catchment.

## Methods

### Study location and sampling sites

We studied ten naturally replicated catchments in eastern Switzerland, with the headwater streams between 2.75 and 5 kilometers in total length (including main and side stems) and running into Lake Constance (covering catchments of in total 115 to 453 hectares). Four streams were located in the less-developed, steeper “Untersee” region to the west, and six were located in the more heavily agricultural, flatter “Obersee” region to the east (Figure 1A). Catchments had varying land use from primarily mixed deciduous and coniferous forest to primarily agriculture, with pockets of higher density housing or industrial uses (Figure 1E).

**Figure 1:**
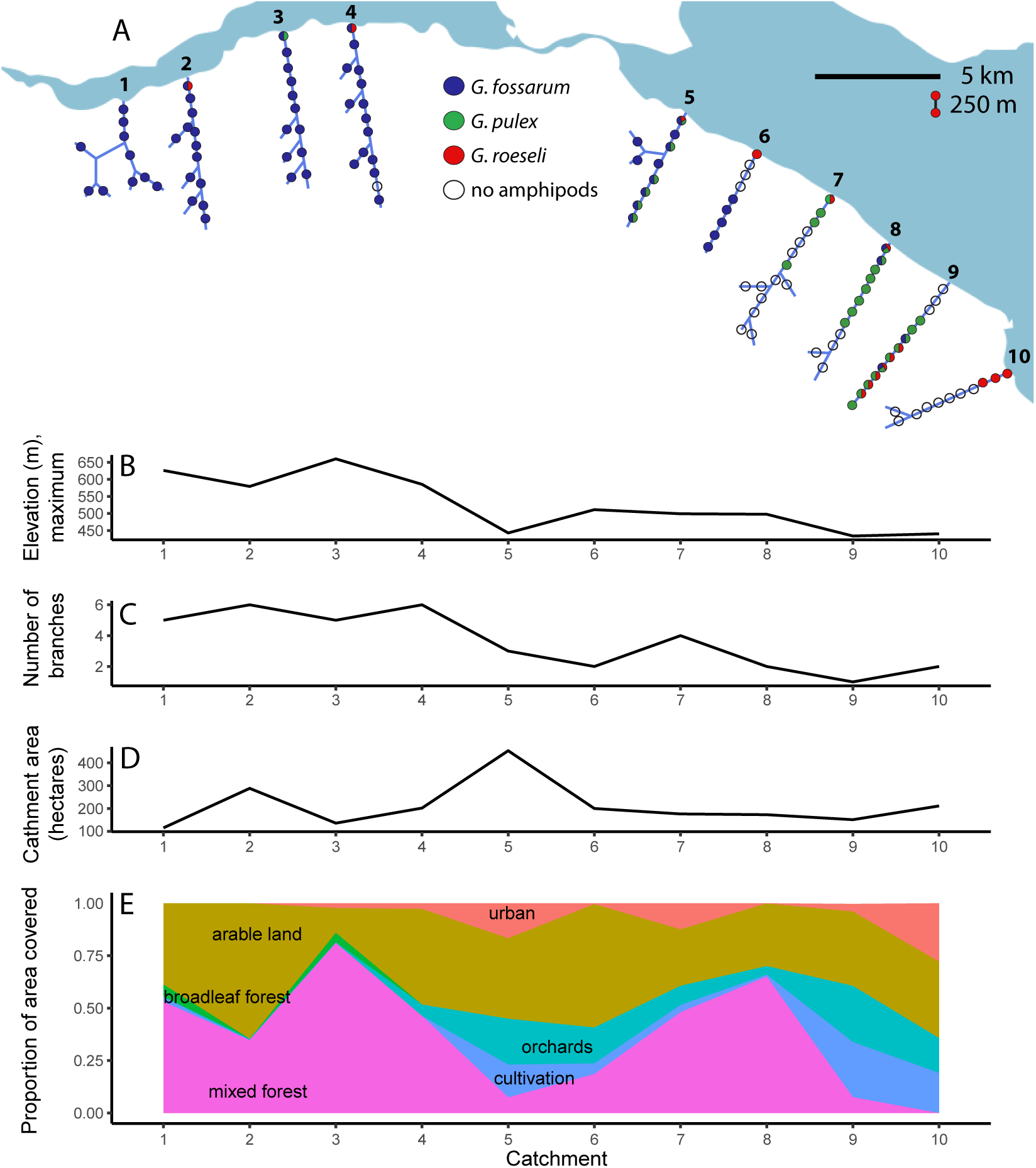
(A) Simplified diagram of 121 sampling points along the branches of ten headwater stream catchments of Lake Constance in Eastern Switzerland. Sampling points are separated by, on average, 250 meters. Colors of sampling points represent whether each species was ever found at the point over a year of sampling effort; divided circles show that multiple species were found at that point, either simultaneously or in sequence. Catchments varied by **(B)** maximum elevation (and thus stream slope), **(C)** branching structure/network complexity, **(D)** catchment area, and **(D)** land use.

In each catchment, streams were divided into 250-meter segments along the main stem.

Side stems less than 450 meters in total length were counted as single segments, while side stems greater than this length were divided into 250-meter segments beginning from the confluence with the main stem. Thus, we sampled throughout the entire network of each stream catchment. A sampling point was established within the segment where habitat and stream flow was as representative as possible of the whole length of the segment. Sampling points were placed as equidistantly as possible given this objective. This resulted in a range of nine to 15 sampling points per catchment, and 121 sampling points in total.

### Data collection

Sampling points were visited four times at roughly three-month intervals, between April/May 2015 and January 2016. Repeat sampling points were within 10 meters of points at the same location. At each visit, three types of data were collected at each sampling point:

1. *Substrate and habitat characterization.* The width of the active channel was measured, and habitat inside a 1 m long section of the stream was visually classified using a 1 × 1 m sampling frame with 0.2 × 0.2 m gridlines. The stream substrate was characterized as area of impermeable surface (bedrock, solidly calcified benthic material, or concrete), rocks >20 cm in diameter, gravel 2.5–20-cm diameter, fine gravel <2.5-cm diameter, sand, mud and clay. The area covered by dead leaves, living terrestrial and large aquatic plants, roots and woody debris, or moss and algae was estimated.
2. *Water chemistry.* A water sample was collected from each sample point and, in the lab, measured for total phosphorus with a spectrophotometer (Varian Cary 50 Bio, Palo Alto, California, USA), and total nitrogen, dissolved organic carbon (DOC), and total organic carbon (TOC), all with a TOC analyzer (Shimadzu TOC-L, Kyoto, Japan).
3. *Amphipod abundance and identity.* After leaf collection, kicknetting was performed across the width of the stream section, such that each type of habitat and substrate type were sampled. Sampling effort was equal per meter of stream width, so that the total time spent kicknetting was greater in wider stream segments. From the kicknet sample, abundance of amphipods was estimated by order of magnitude: 0, 1–10, 10–100, 100–1000, or more than 1000. From each sample roughly 30 amphipods were collected and preserved in alcohol, or all of the amphipods present if there were less than this number. The collected amphipods were chosen to represent the range of sizes present in the sample, but not including those which were too small to reliably identify to species. After being brought to the lab, amphipods were identified to species.

### Land-use data

All spatial analysis was done using ArcGIS 10.2.2 (ESRI, Redlands, California, USA). Spatial information about streams was extracted from the Swiss national 1:25,000 scale water network (Swisstopo 2007). This was overlaid on a digital elevation model accurate to within two meters (Swisstopo 2003). From this, the elevation of each sampling point and its upstream distance from the outlet on Lake Constance were calculated.

Land cover within the catchments was classified using a combination of data sources. We used as the basis the CORINE land cover (2012) European Environment Agency (EEA) land-use classification (Bossard *et al.* 2000), produced from Indian Remote Sensing (IRS) P6 LISS III and RapidEye imagery with a Minimal Mapping Unit of 25 hectares and positional accuracy of, at a minimum, 100 meters. To add specificity to CORINE’s agricultural classification and because orchards often have higher herbicide application, we added the area of vine and orchard fruit cultivation from the Swiss national 1:25,000 scale vector map (Swisstopo 2010). After this distinction, land cover within the catchments fell into nine categories: discontinuous urban fabric, industrial or commercial units, non-irrigated arable land, complex cultivation patterns, fruit orchards and vine cultivation, broad-leaved forest, mixed forest, inland marshes, and water bodies. The area of land falling into each land use category was calculated for each study catchment in total, as well as for a 50-meter radius circular area surrounding each individual sampling point.

### Statistical analysis

All statistical analyses were performed in R version 3.3.2 (R Core Team 2016). The presence or absence of amphipod species was examined using the ‘HMSC’ package for Hierarchical Modeling of Species Communities (Ovaskainen *et al.* 2017), a Bayesian joint species distribution model. We incorporated environmental covariates and the sampling structure, with catchment and sampling point as random factors representing spatial context. We used the default priors of the package and modeled species occurrences using the Bernoulli distribution and a probit link function (additional information on model specification in Appendix I). MCMC chains were run to 100,000 iterations, with a burn-in period of 1,000 iterations and subsequently thinned to include only every 100^th^ sample of the posterior distributions. We primarily compared two models. The first included only the spatial random effects (“S”). The second included three types of factors: spatial random effects, prior amphipod occurrence, and environmental covariates (“SPE”). To use the information about which amphipod species had been present at a point previously, these two models were made using data only from the second through fourth sampling timepoints (at the first sampling timepoint, there was no prior presence-absence information). For comparison purposes, we repeated the model with random effects plus environmental factors, but not including the data about prior amphipod occurrence, with the complete dataset of all four timepoints (“SE_Full_”). We ran two additional models – one with spatial random effects plus information on the prior presence of amphipod species (“SP”), and another with the random effects plus all other environmental covariates described above (both field-measured and GIS-derived; “SE”) – the results of which are presented in Appendix I.

Overall model fit was assessed using Tjur’s R^2^ (Tjur 2009), which is defined as the difference between the mean fitted value of sampling units where species are present and the mean fitted values where species are absent. Importance of environmental covariates was assessed in two ways. First, parameter estimates of the association between environmental covariates and presence/absence of individual species were extracted as 95% central credible intervals of the posterior sampling distribution. Where this central credible interval did not overlap with zero, the particular environmental covariate was deemed to have a strong directional association with the presence or absence of the species in question. Then, the explained variance in presence/absence of each species was partitioned among all explanatory variables, which were grouped for presentation into broad categories, as well as to random effects at the both sampling scales (catchment and sample point). Finally, we assessed the potential co-occurrence of species, or “hypothetical species association patterns” (Aivelo & Norberg 2017), by extracting the residual associations between species from the latent part of the model framework. A positive residual association indicates that species occur together more frequently than would be predicted by their calculated niches, while a negative residual association indicates that their niches would predict them to co-occur more frequently than they do in practice. These putative species associations are depicted by the median value of posterior samples and expressed as correlations.

## Results

### Spatial and temporal patterns in distribution

Our sampling revealed a pronounced spatial pattern in species distributions, with *Gammarus fossarum* the only species present upstream of outlets in the western catchments and three different species (*G. fossarum*, *G. pulex*, and *G. roeseli*) present in eastern catchments, but rarely coexisting (Figure 1). Of these three, *G. fossarum* and *G. pulex* are native species while *G. roeseli* is non-native but considered naturalized since it arrived in the 1800s. Across the whole study region and sampling year, mean species richness at outlet points was 1.25 species (range 0– 3), and at non-outlet points was 0.69 species (range 0–2) at points that had water (some stream reaches were dry at some sampling times, see below). No non-outlet point ever had three species present. From this, we concluded that outlets were more representative of the lake’s species pool of five to six species (Altermatt *et al.* 2014, 2016) than of stream communities, and we excluded outlet points from subsequent analyses.

Site occupancy was fairly stable through time, with no change in species composition at a sampling point in 78% of the possible transitions from one timepoint to the next. There were few changes from single-species to multi-species occupancy (3% of possible transitions) or vice versa (2% of possible transitions). There was also one change from a point being occupied by one to a different species (0.3% of transitions; Figure 2). The most common change at the sampling point level was from being occupied to being unoccupied (11% of possible transitions), in large part due to seasonal drying of some stream reaches. Few (5/17) of these dried stream reaches were reoccupied, and in all of these cases they were reoccupied by the same species which had been present before the drying event. In one of the five cases, an additional species co-established at the re-wetted stream reach. Overall, this shows that there is nearly zero turnover in species dominance amongst occupied sampling points and little chance for “new” species to establish after disturbance has rendered some patches unoccupied.

**Figure 2:**
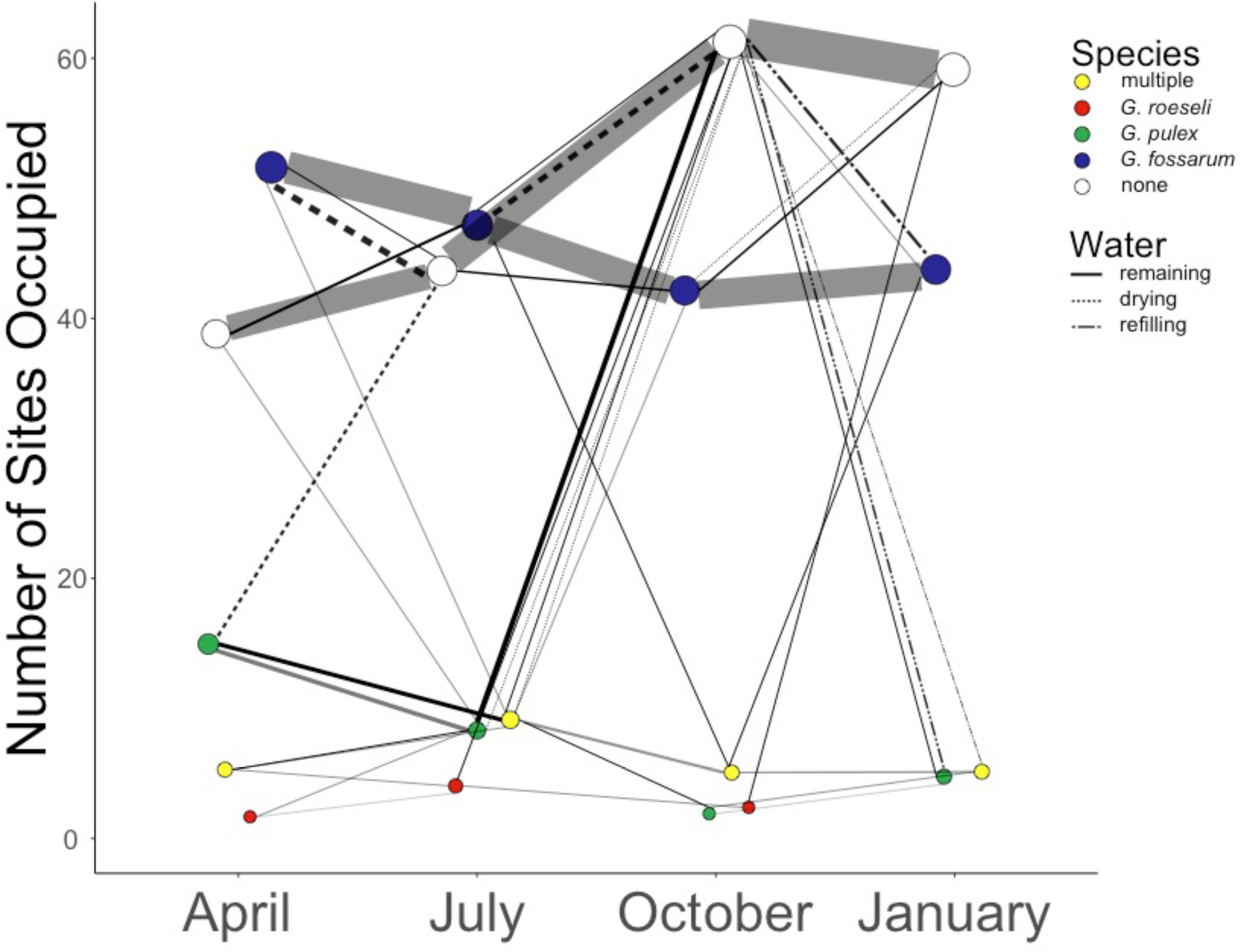
Maintenance (gray lines) and changes in occupancy (black lines) at the 111 sampling points (outlets were excluded in this analysis) over a year of visits. Size of circles is scaled to the proportion of points in this state, and line thickness is scaled to the number of state transitions of this type over the given sampling interval. Less than ten sampling points contained multiple coexisting species (yellow points) at any sampling visit; the overall proportion of points containing zero species (white circles) increased over the duration of the study, due in large part to sites which dried out completely (dotted lines). Only some of these intermittent reaches were recolonized when water returned in the final sampling visit (dash-dotted lines).

### Comparing joint species distribution models

The spatial arrangement of the sampling points was shown to be important in explaining the presence and absence of different species through several different metrics. The *S* model using only the random effects of catchment and sampling point for the second through fourth timepoints (n=390) explained 51% of the variation in species presence and absence. The full “SPE” model (n=237) explained 71% of variation. The models with either prior amphipod occurrence or environmental covariates separately had intermediate model fits (Appendix I). By comparison, the *SE*_*Full*_ model across all sampling timepoints but using only spatial arrangement and environmental covariates (n=367) explained 64% of variation. This suggests that both environmental data and data about species distributions at prior sampling times are important and do not convey the same information.

### Abiotic influences on species distribution

In the *SPE* model, only a few variables had strong directional effects (defined using the 95% central credible interval of the posterior distribution of the association) on the presence or absence of amphipod species (Table 1). Despite their strong directional effects, these variables did not necessarily account for a large proportion of the variance in occurrence patterns (Figure 3); for instance, the association between the area of substrate covered by leaves and the occurrence of *G. pulex* accounted for only 1.7% of the explained variation in the species’ occurrence. For *G. pulex*, the area of the streambed covered by leaf litter, the proportion of area surrounding a point made up of arable land, and the previous presence of *G. pulex* were strong predictors. For *G. fossarum*, latitude, the proportion of catchment area covered by orchards, the dissolved organic carbon in the water, and the previous presence of *G. fossarum* were strong predictors. And for *G. roeseli*, the proportion of area surrounding a point used for industrial or commercial purposes and the previous presence of *G. pulex* were strong predictors. Other important factors in the *SE*_*Full*_ model, such as the association between *G. fossarum* and previous drying or the area of moss and algae on the substrate, no longer had strong directional effects when both environment and previous species occurrence information were integrated into the same model (Table 1). Although other factors measured at the sampling point or catchment level did not have strongly directional effects, they nevertheless contributed greatly to explaining the variation in species occurrences when a variance partitioning was conducted on the *SPE* model (Figure 3). For example, land use in the catchment accounted for 32% of the explained variation in the occurrence of *G. fossarum*, and 6% of the explained variation in *G. pulex*; while variables measured at the point level which did not have strong directional effects nevertheless combined to account for 24% of the explained variation in the occurrence of *G. fossarum*, 13% of explained variation in *G. pulex*, and 5% of the explained variation in *G. roeseli.* The density of posterior distributions of all associations between measured variables and species occurrences are presented in Figure S1.

**Table 1.** Proportion of variance explained by factors with strong directional effects (+ or -) on the occurrence of species in two hierarchical joint species distribution models: that including spatial random effects, previous occurrence, and environmental covariates (*SPE*), calculated over the last three sampling timepoints because information on prior amphipod occurrence was not available at the first sampling timepoint; or spatial random effects plus environmental covariates considered over all four sampling timepoints (*SE_Full_*). Only factors having this strong effect (defined as the 95% central credible interval of the association between the factor and the species presence/absence being non-overlapping with zero) for the given species in at least one of the models (models are described in the methods section) are included in the table. Factors without a strong directional effect for a given model are indicated with “n.s.”, and factors not included in a given model are indicated by “N/A”. The random effects associated with catchment and sampling point are included for reference for all species and models and highlighted in gray to differentiate them from the effects of measured covariates.

|  | *SPE* model | *SE<sub>Full</sub>* model |
| --- | --- | --- |
| **(A) *G. pulex*** |  |  |
| Catchment | 0.501 | 0.445 |
| Point | 0.002 | 0.003 |
| *G. pulex* previous presence (+) | 0.020 | N/A |
| Arable land at point (+) | 0.013 | 0.015 |
| Area of leaves (-) | 0.012 | 0.017 |
| **(B) *G. fossarum*** |  |  |
| Catchment | 0.001 | 0.026 |
| Point | 0.002 | 0.074 |
| Latitude (+) | 0.112 | 0.135 |
| *G. fossarum* previous presence (+) | 0.075 | N/A |
| Orchard in catchment (+) | 0.067 | n.s. |
| Dissolved organic carbon (-) | 0.056 | 0.023 |
| Previous drying (-) | n.s. | 0.018 |
| Area of moss, algae, macrophytes (-) | n.s. | 0.017 |
| **(C) *G. roeseli*** |  |  |
| Catchment | 0.634 | 0.566 |
| Point | 0.001 | 0.001 |
| *G. pulex* previous presence (+) | 0.004 | N/A |
| Industrial/commercial at point (+) | 0.003 | 0.003 |

**Figure 3:**
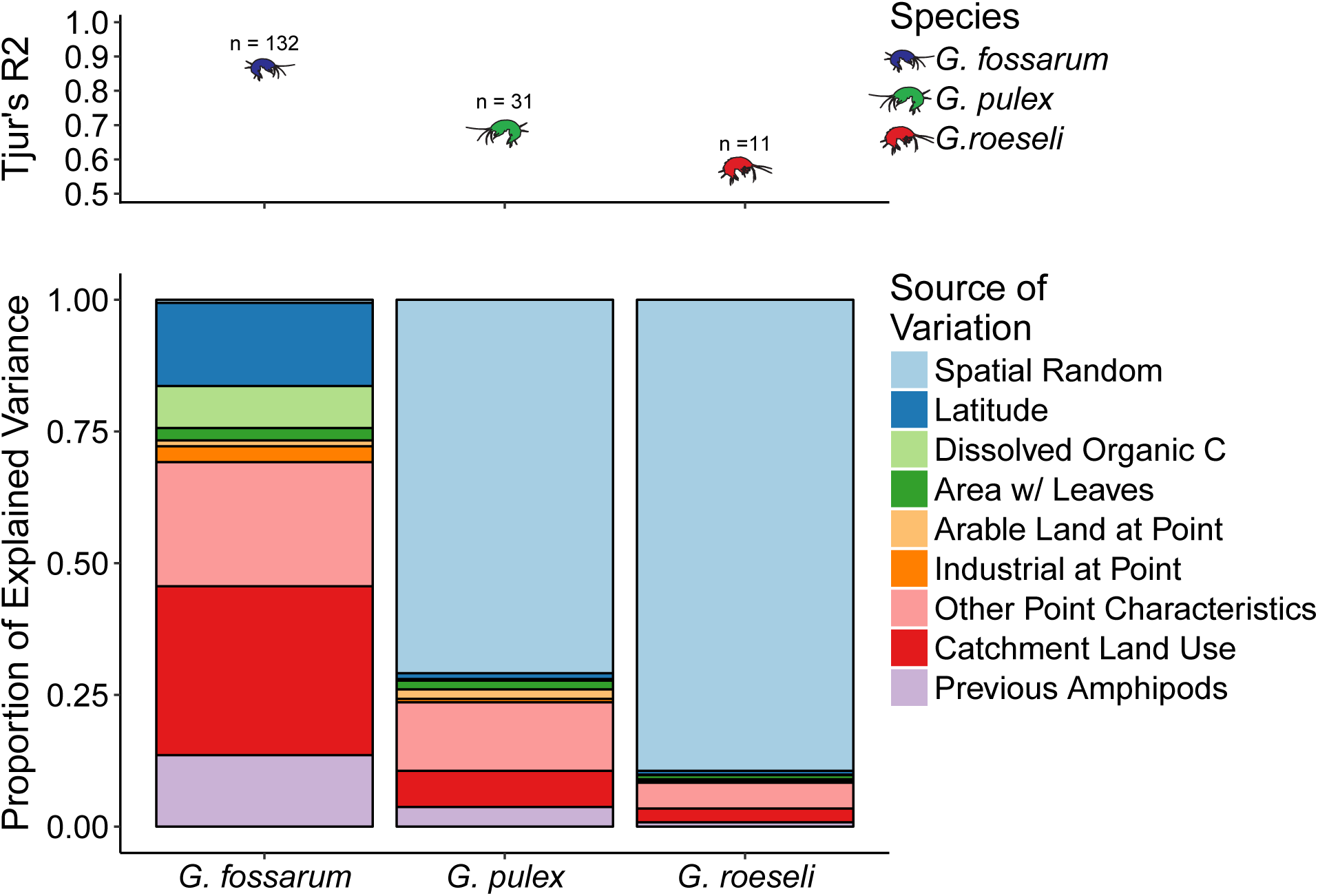
(A) The amount of variance in species occurrences (Tjur’s R^2^) explained by model components for each species individually, and the number of times each species was detected in the field sampling. **(B)** Variance partitioning of factors used in the *SPE* model (spatial random effects, prior amphipod occurrence, environmental covariates) in relation to presence/absence of each species found in the study. Stacked bars show the proportion of the total explained variance, indicated in panel (A) for each species; overall, 71% of total variance in the dataset was explained by the model.

### Co-occurrence of amphipod species

After accounting for the influence of these factors, there were putative species associations between different amphipod species in the *SPE* model: weak positive associations at the catchment level, and strong positive and negative associations at the sampling point level (Figure 4). At the sampling point level, *G. fossarum* rarely co-occurred with either of the other species despite somewhat-similar habitat requirements (median residual correlation = -0.74 to *G. pulex* and -0.75 to *G. roeseli*). Conversely, *G. pulex* and *G. roeseli* co-occurred much more frequently (median residual correlation = 0.99) than would have been predicted either by random chance or based on their individual habitat requirements. At the catchment level, pairs of species co-occurred slightly more frequently (residual correlations of 0.10–0.25) than would have been predicted either by random chance or by the niches constructed based on our measured factors (spatial arrangement, previous species occurrence, and environmental covariates).

**Figure 4:**
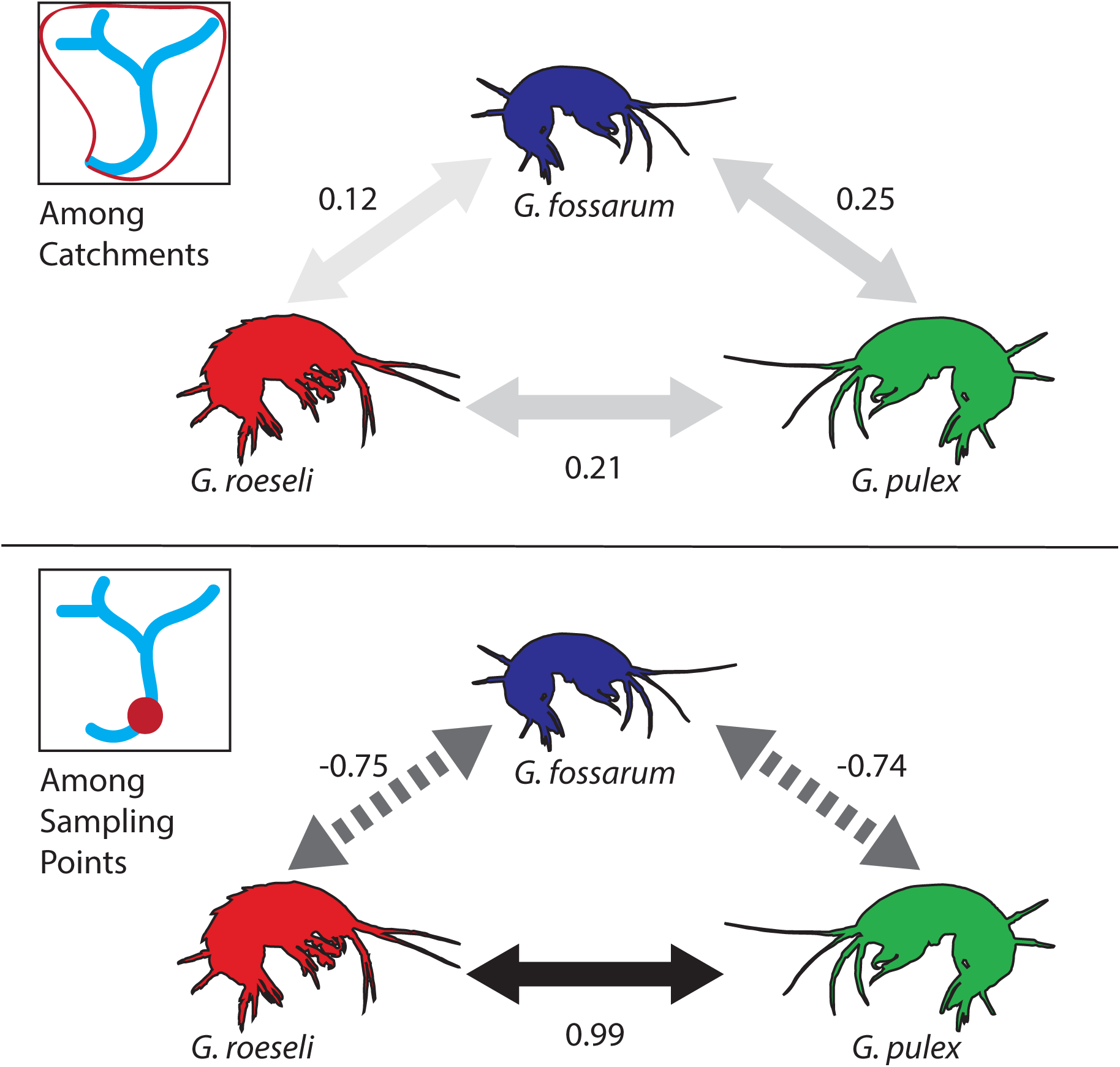
Residual associations in occurrence between different species at the catchment and sampling point level, after the influence of spatial relationships, environmental covariates and previous amphipod site occupancy have been taken into account by the *SPE* model. Arrows are colored by the strength of the association; positive associations (represented by solid lines) indicate that species co-occur more frequently than would be predicted by environmental covariates, spatial random effects, and previous amphipod site occupancy, while negative associations (represented by dashed lines) indicate that species co-occur less frequently than predicted.

## Discussion

It has been commonly observed that species do not always co-exist where might be expected. We examined the distribution of three locally-dominant and occasionally co-occurring amphipod species in order to disentangle the ecological processes behind their occupancy patterns in stream catchments. As expected, the environmental factors we measured explain part of the variation in these species’ distribution between and within catchments. Overall, however, individual environmental factors rarely had strongly positive or negative effects on species occurrences, and a large amount of variance remained unexplained by environmental variables, as is commonly found (Cottenie 2005; Ozinga *et al.* 2005; Heino *et al.* 2015). Importantly, these three species are not dispersal-limited at the scale of our studied headwater stream catchments (Altermatt et al. 2016), ruling out another common mechanism shaping community composition.

Using a joint species distribution modeling approach, we show that this unexplained variance can be assigned to putative species interactions. While much experimental work on community assembly and species interactions has been done in plant communities -- where individuals are immobile, order of arrival can be easily manipulated and neighbors may be removed from a community (i.e., Choler *et al.* 2001; Körner *et al.* 2008) -- this is more challenging when working with animals in flowing-water systems. We used an analytical approach which allowed us to infer species interactions from observational data (Aivelo & Norberg 2017; Ovaskainen *et al.* 2017) without performing manipulations. In past studies, competition has been assumed to be the primary species interaction shaping amphipod communities: for example, *G. pulex* rarely co-occurred with another sympatric species, *G. duebeni*, in rivers in France, and this was attributed to hypothesized strong competition between the two species (Pinkster *et al.* 1970). However, in no amphipod distribution studies that we are award of has competition been directly measured or indirectly inferred, and it was rather proposed as an overarching mechanism. Our results now suggest a more nuanced role of competition.

After accounting for important environmental factors we found strong negative species interactions, but because different species are dominant in different catchments, we ruled out that one species has an absolute fitness and competitive advantage over the others; in this case, the same species should have dominated all of our catchments. Which species dominated which catchment was also not satisfactorily explained by environmental variables, suggesting that the identity of the “winner” is also not deterministically driven by niche differences. This conclusion is based on the fact that at the catchment scale, we found that species coexisted more frequently than would be expected based on environmental factors. In particular, in two catchments where multiple species co-occurred throughout the length of the stream, they did so at roughly equal densities over the course of the entire study period (Table S2). These two catchments were not particularly close to each other and had some quite different land use (Figure 1) and other habitat characteristics (Supplementary Data), making it highly improbable that some particular abiotic variable promoted coexistence. The ability of the species to coexist was also found in laboratory experiments, where *G. fossarum* and *G. roeseli* each had equal (~90%) survival over short-term experiments, regardless of whether maintained separately or together in mesocosms at equal densities (Little et al., in review). This rules out strong, frequency-independent competition between the species, to the degree that would cause competitive exclusion when the species are at similar densities. And yet most strikingly, at the scale of sampling reaches we found a putative negative species association between the two most common species in the region, *G. fossarum*and *G. pulex*. Indeed, classic studies in France (Pinkster *et al.* 1970; Piscart *et al.* 2007) and Britain (Hynes 1954) found that derived environmental preferences (i.e., niche differences) were insufficient in explaining the distribution of freshwater amphipods. Instead, in our data the previous occurrence of the same species at a sampling point was the only strong positive predictor of the occurrence of *G. fossarum* and *G. pulex*. Thus, it is clear that coexistence of species depends on scale (Hart *et al.* 2017), and we indeed saw coexistence at the catchment but only rarely at the reach scale.

Consequently, the remaining question is what the source of these differing patterns of coexistence is, and how the strong negative association between the two most common species is shaped. Neither a pure environmental filtering nor competitive exclusion perspective offer convincing explanations in our analysis. Alternatively, priority effects are thought to be common in ecosystems (Alford & Wilbur 1985; Almany 2003; De Meester *et al.* 2016). They are, however, generally difficult to quantify through observational study because the history of community assembly is rarely known (Fukami 2015). Several patch characteristics are associated with promoting priority effects among functionally-similar species. These mechanisms typically allow early-arriving species to quickly grow to large population sizes: for instance, small patch size and a stable environment with high resource supplies and/or lack of predation (Fukami 2015). In linear habitat networks such as streams, which are surrounded by an unsuitable (terrestrial) habitat matrix and where each habitat patch (stream reach) is connected to only a very small number of other patches, priority effects may play an outsized role due to spatial blocking. Notably, after a large-scale disturbance such as a flood, drought, or a chemical spill, purely aquatic animals primarily colonize stream networks from the outlet up. Thus, if a species first colonizes a stream reach near the outlets, this species encounters low resistance while dispersing further upstream and may quickly rise to high densities in these patches as well. Conversely, it may become very difficult for another newly-arriving species to pass through these initial downstream habitat patches en route to suitable (potentially even empty) upstream reaches, once a prior species is present. This may explain why although we did see disturbance in our study streams – some dried out during the summer drought, but primarily in upstream reaches – these streams were not recolonized by new species, as they likely did not have access to them.

Similarly, distributions in a partially overlapping set of streams measured at a coarser spatial scale did not change much over two years (Altermatt *et al.* 2016). We assume that after events such as heavy pesticide application to surrounding farm fields, species turnover in a catchment could occur (Thompson *et al.* 2016) as long as the disturbance reached all the way downstream to the lake and provided access for all species in the regional species pool. Priority effects have been invoked to explain macroinvertebrate community composition at individual stream or river reaches (McAuliffe 1984; Palmer *et al.* 1996), but as far as we are aware, the role of priority effects in excluding species at the catchment or network level has not yet been investigated in natural riverine systems.

There are several other specific mechanisms supporting and consistent with the role of priority effects in structuring these amphipod communities. First, intraguild predation is one setting where order of arrival to a habitat patch may be particularly important in determining community structure in many ecosystems (for example, Blaustein & Margalit 1996 in temporary ponds). And indeed, intraguild predation has been reported to be common in various *Gammarus* species pairs, often at a stronger intensity by one species than the other (Macneil *et al.* 1997).

This mechanism was used to explain how a non-native species can extirpate a less aggressive native species from its home range, but would almost certainly prevent the weaker species from establishing a stable coexisting population amongst the stronger species were the tables reversed. Secondly, mate limitation may also prevent new species from moving into a catchment dominated by a single other species. *Gammarus* species have been shown to have varying abilities to differentiate between potential mates of different species (Kolding 1986; Dick & Elwood 1992). Some form interspecific copulatory pairs even when mates of their own species are available, and others do so only if mates of their own species are unavailable; these pairings lead to the fertilization of eggs, but offspring fail to develop (Kolding 1986). Generally, when at a low density, males may mate with females of the dominant species and thus fail to sustain their own population.

### Conclusion

We found that although part of the variation in the distribution of *G. fossarum* could be explained by environmental measures, our putative negative species interactions were highly important in explaining the distinct spatial patterns we observed. Multiple species rarely coexisted, even in reaches that would seem to be suitable for more than one species. This leads to a classic problem: despite knowledge of environmental conditions, it can be difficult to predict where a given species will be found if other factors are preventing it from occupying all suitable niche space (Hutchinson 1957). Competition is often invoked as a probable cause for one species to exclude another. Yet we show that even species that can coexist while competing in some circumstances, do not do so in all conditions. Order of arrival and initial establishment of a robust population may determine which of several competitors occupy a community. Furthermore, river networks represent a unique spatial setting for such considerations, since colonization by aquatic organisms is in many cases directional (downstream to upstream or vice versa). For example, established dominant species in downstream reaches have a head start towards colonizing empty upstream patches and may prevent newly-arriving species from passing through occupied habitat patches to reach empty ones. When the species of interest are dominant species, which have strong effects on community structure and ecosystem function, this may be especially problematic: if niche preemption prevents species from coexisting, it will be difficult to infer levels of ecosystem functioning.

## Acknowledgments

The authors would like to thank the Kanton Thurgau Office of the Environment for helping ensure easy access to sampling sites, and all landowners whose property we crossed. We are also indebted to Pravin Ganesanandamoorthy, Elvira Mächler, Simon Flückiger, and Katharina Kaelin for help with fieldwork, and the Eawag AuA laboratory for chemical analysis of water samples. Tadashi Fukami, Marcel Holyoak, Brad Taylor, and Hanna Kokko provided helpful feedback while the manuscript was in preparation. This project was funded by Swiss National Science Foundation grant PP00P3_150698.

